# Effects of Age on Cross-Cultural Differences in the Neural Correlates of Memory Retrieval

**DOI:** 10.1101/2024.04.25.591227

**Authors:** Isu Cho, Krystal R. Leger, Ioannis Valoumas, Ross W. Mair, Joshua Oon Soo Goh, Angela Gutchess

## Abstract

Culture can shape memory, but little research investigates age effects. The present study examines the neural correlates of memory retrieval for old, new, and similar lures in younger and older Americans and Taiwanese. Results show that age and culture impact discrimination of old from new items. Taiwanese performed worse than Americans, with age effects more pronounced for Taiwanese. Americans activated the hippocampus for new more than old items, but pattern of activity for the conditions did not differ for Taiwanese, nor did it interact with age. The engagement of left inferior frontal gyrus (LIFG) differed across cultures. Patterns of greater activity for old (for Americans) or new (for Taiwanese) items were eliminated with age, particularly for older Americans. The results are interpreted as reflecting cultural differences in orientation to novelty vs. familiarity for younger, but not older, adults, with the LIFG supporting interference resolution at retrieval. Support is not as strong for cultural differences in pattern separation processes. Although Americans had higher levels of memory discrimination than Taiwanese and engaged the LIFG for correct rejections more than false alarms, the patterns of behavior and neural activity did not interact with culture and age. Neither culture nor age impacted hippocampal activity, which is surprising given the region’s role in pattern separation. The findings suggest ways in which cultural life experiences and concomitant information processing strategies can contribute to consistent effects of age across cultures or contribute to different trajectories with age in terms of memory.

## Effects of Age on Cross-Cultural Differences in the Neural Correlates of Memory Retrieval

What and how we remember is influenced by various individual difference factors, including the experiences of living in specific cultural contexts. Although research has revealed ways in which culture influences cognitive processes and their corresponding neural correlates, very little work has investigated the effects of culture with age. Thus far, some of the findings show consistency across cultures, with age effects generalizing across cultures or cultural differences emerging across younger and older adults. For example, the tendency for higher rates of memory errors with age, primarily documented in Western populations (Balota et al., 1999; Roediger & McDermott, 1995), occurs for Turkish and American older adults (Gutchess & Boduroglu, 2019). Reductions in source memory (Chua, Chen, & Park, 2006) and item-context binding (Yang, Li et al., 2013) with age generalize across cultures. Cross-cultural differences in the *types* of memory errors (Schwartz, Boduroglu, & Gutchess, 2014; see also Wang et al., 2019; 2021) extend to older adults (Gutchess & Boduroglu, 2019). Other research, however, suggests ways in which aging processes unfold differently across cultures. For example, American older adults employ a clustering strategy in memory more than Chinese older adults (Gutchess et al., 2006) and Western advantages for categorical information may exaggerate memory differences with age across cultures (Yang, Chen, Ng, & Fu, 2013). Memory strategies focused on the self are less effective for Taiwanese older adults, compared to Americans older adults (Zhang et al., 2020). Taken together, the behavioral memory studies show evidence for both the consistency of age-related changes across cultures and cultural differences that are more pronounced with age.

Despite the growing number of studies investigating the influence of culture on memory with age, there has been virtually no research incorporating measures of brain structure and function. There are only two studies (Chee, Zheng, Goh, Park, & Sutton, 2011; Goh et al., 2007) that directly compare younger and older adults across cultures with a cognitive neuroscience approach. Specifically, Chee et al. (2011) compared brain structure (i.e., cortical thickness and volume) in younger and older East Asians and Westerners to test whether brain structure differs across cultures given the previous evidence of cross-cultural differences in cognitive processes (i.e., East Asians: holistic view vs. Westerners: analytic view; Nisbett & Miyamoto, 2005). They found that the cultural difference in brain structure in younger adults was reduced in older adults in general and that the age-related structural differences were pronounced and consistent across cultures. Similarly, the sole study using functional imaging (Goh et al., 2007) investigated the effects of age and culture on visual processing, and results showed some universal effects of age on hippocampal activity during object-scene binding. In terms of cultural differences, activity in object processing regions was impacted by age in Easterners more than Westerners. Importantly, this study used a passive viewing task that did not directly probe memory performance. Although the two neuroimaging studies revealed consistent effects of age across culture, as well as attenuated cultural differences with age, it is still unknown how culture and age interact with each other on memory processes and their neural correlates.

The present study investigates cross-cultural differences in memory with age, comparing younger and older adults from Western (US) and Eastern (Taiwan) cultures using behavioral and neuroimaging methods. Past behavioral research comparing younger adults across cultures demonstrates that Americans can have more accurate memory for the features of objects compared to East Asians (e.g., discriminating one particular bicycle in memory from another visually similar exemplar), indicating cultural differences in memory specificity. The cultural differences in memory for objects have been observed regardless of whether objects were presented alone or against a background (Millar, Serbun, Vadalia, & Gutchess, 2013) and regardless of the congruency of the object-background pairing (Mickley Steinmetz, Sturkie, Rochester, Liu, & Gutchess, 2018). In a more recent study (Leger et al., 2024b), American’s higher level of memory specificity compared to East Asians was found not only for concrete everyday objects but also for abstract figures. These results converge with related work on cultural differences in the amount of detail contained in autobiographical memories (e.g., Wang, Hou, Tang, & Wiprovnick, 2011; see Wang, 2021 for a review). In some cases, cross-cultural differences in memory even extend to the discrimination of old from new items (Leger & Gutchess, 2021; Leger et al., 2024b), with Americans exhibiting higher levels of memory performance compared to East Asians. These findings indicate powerful cultural differences in memory across different contexts and types of stimuli.

Despite the number of demonstrations of cross-cultural differences in memory specificity and even old-new differences in memory, how the effects of age and culture interact is underexplored. Establishing to what extent memory impairments are a pervasive effect of age or whether they vary depending on environmental factors or learned strategies based on cultural context is an important question. Impairments in memory, particularly in terms of the level of detail or specificity, can be pronounced with age and in age-related disorders such as amnestic mild cognitive impairment (aMCI) or Alzheimer’s disease (AD) (Balota et al., 1999, Bowman et al., 2019; Paige, Cassidy, Schacter, & Gutchess, 2016; Gellersen, McMaster, Abdurahman, & Simons, 2024), so exploring the extent to which environmental factors can modulate the effects of aging on memory could inform the development of effective interventions to mitigate age-related deficits in memory.

Pattern separation, creating a new neural code for an item that differs from one seen previously, is one mechanism that has been proposed to support accurate and detailed memory (Yassa & Stark, 2011). This process contrasts pattern completion, which relies on reactivating the same neural code as for an exemplar seen previously. Pattern separation supports mnemonic discrimination or distinguishing similar experiences from each other. For example, knowing the perceptual features that distinguish one’s sneakers from others’ sneakers allows one to retrieve the correct footwear at the end of yoga class. Regions of the hippocampus exhibit distinct responses when pattern separation vs. completion is required (Baker et al., 2016; Bakker, Kirwan, Miller, & Stark, 2008; Doxey & Kirwan, 2015; Lacy, Yassa, Stark, Muftuler, & Stark, 2011; Liu, Gould, Coulson, Ward, & Howard, 2016; Yassa & Stark, 2011; but see Quiroga, 2020). As reviewed by Leal & Yassa (2018), the Cornu Ammonis 1 (CA1) region exhibits a linearly decreasing response to interference between similar representations in memory, with the strongest signal when there is no interference. In contrast, the dentate gyrus (DG) and CA3 support pattern separation, depending on the input (these regions are referred to as DG/CA3, as they cannot be distinguished from each other in *in vivo* human brain images). For example, the activity in the DG/CA3 is sensitive to subtle changes in items (Lacy et al., 2011) and an individual with lesions to the DG had difficulty in identifying similar items (i.e., lures) (Baker et al., 2016). With age, there is evidence for impairment of pattern separation, behaviorally and neurally (Leal, Noche, Murray, & Yassa, 2017; Reagh et al. 2018; Stark, Stevenson, Wu, Rutledge, & Stark, 2015; Stark, Yassa, Lacy, & Stark, 2013; Stark, Yassa, & Stark, 2010; Yassa, Mattfeld, Stark, & Stark, 2011). These impairments are further exaggerated in aMCI (Stark et al., 2013) and related to markers of AD, such as beta amyloid (Adams et al., 2022). Tasks testing the ability to discriminate old from similar and new items, such as the Mnemonic Similarity Task (MST; Stark, Kirwan, & Stark, 2019), are sensitive at detecting memory impairments with age even before individuals have reached clinical levels of impairment (Stark et al., 2013). This contrasts traditional measures of old/new recognition memory that cannot distinguish cognitively unimpaired from impaired older adults.

For the hippocampus, the structure implicated in pattern separation and in forming and retrieving detailed memory representations, there is some evidence for cultural differences. Compared to Americans, East Asians had greater activation in the left hippocampus, along with the left parahippocampal and left fusiform gyri, when they successfully formed detailed memory representations during the encoding stage that supported accurate performance at test (i.e., subsequent memory design; Paige, Ksander, Johndro, & Gutchess, 2017). However, initial tests of pattern separation as a candidate mechanism did not provide support for the idea that cultural groups differ in this mechanism. Behaviorally, cultural differences emerged more broadly than merely for lures, with Americans exhibiting higher levels of memory accuracy for differentiating pictures of target objects from similar lures as well as from new objects, compared to East Asians (Leger & Gutchess, 2021; see also Leger et al., 2024b). Neural measures, which provide a stronger test of the involvement of the hippocampus in pattern separation, also failed to reveal cultural differences in the engagement of hippocampal regions during the discrimination of correct rejections from false alarms for similar lures during retrieval (Leger et al., 2024a). Note, however, that this study did not use high-resolution imaging to target the precise regions implicated in pattern separation and completion (e.g., DG/CA3; CA1). Nevertheless, testing for potential cultural differences in pattern separation and associated neural regions in older adult samples should provide a stronger test than the prior studies with younger adults, given the changes to these processes and brain regions that can occur with age.

In addition to testing the joint effects of age and culture on pattern separation, the present study will also investigate the effects of these factors on old/new discrimination in memory. Although this measure is not precise in targeting age-related memory decline related to aMCI or AD, cultural differences of interest have emerged for old vs. new comparisons in memory (Leger & Gutchess, 2021; Leger et al., 2024b). Discriminating foils (new items) from targets (old items) during retrieval was associated with different patterns of recruitment of left inferior frontal gyrus (LIFG), left middle frontal gyrus, and right hippocampus across culture groups (Leger et al., 2024a). Interestingly, Americans tended to engage these regions more when correctly identifying new compared to old items, whereas Taiwanese engaged the regions more for old than new items. We speculated that these differences could reflect cultural differences in memory states (Long & Kuhl, 2021), such as attending to old (retrieval orientation) vs. new (encoding orientation) information, which could support more detailed encoding processes for Americans such that they experienced less interference at retrieval. In contrast, if Taiwanese do not encode information into memory in as much detail, they may be prone to experience more interference at retrieval, necessitating the engagement of regions such as LIFG to resolve interference (e.g., Badre, 2008; Badre & D’Esposito, 2007; Badre & Wagner, 2007). Although these proposed explanations require direct tests using appropriate tasks that manipulate interference, the findings provided a basis to compare groups of older adults across cultures on old/new discrimination during memory retrieval.

The present study builds on the younger adult samples investigated in Leger et al. (2024a) by adding older adult samples of Americans and Taiwanese. To this end, younger and older American and Taiwanese participants completed the MST, a surprise memory recognition task for targets (i.e., studied objects), lures (i.e., objects similar to the targets), and foils (i.e., new objects) during fMRI scanning. To assess the effects of age on cultural differences in memory, we compared the groups on memory decisions related to pattern separation (i.e., correct rejections vs. false alarms for lures) and old vs. new discrimination (i.e., hits for targets vs. correct rejections of foils). Relating to age, we hypothesized that one of two potential patterns of outcomes may occur. For one, cultural differences in memory specificity could be enlarged in older adults compared to younger adults. Such a pattern would reflect the greater accumulation of culturally-specific lifetime experiences with age (Gutchess & Cho, 2024; Gutchess & Gilliam, 2022; Park, Hedden, & Nisbet, 1999). Any potential buffering effects of culture may be even more pronounced with age in the case of a cognitively demanding task, such as memory discrimination. For the samples of younger adults in the present study, no cultural differences in behavioral performance were observed in Leger et al. (2024a) (perhaps reflecting the visually-impoverished environment of the scanner compared to the lab). For older adults, therefore, we predicted cultural differences in memory performance such that Americans would have higher levels of memory discrimination performance than Taiwanese. Neurally, we predicted cultural differences in the hippocampus, given its role in pattern separation which declines with age, and in LIFG, as seen for younger adults (Leger et at. 2024a). Age-related changes in memory could exaggerate cultural differences in older adults, as they may deploy culturally-preferred strategies to compensate for age-related declines. The second possibility is that memory changes with age may be pervasive and potentially universal, reflecting the strong influence of biological aging processes in memory (Head, Rodrigue, Kennedy, & Raz, 2008; Korkki, Richter, Jeyarathnarajah, & Simons, 2020; Yassa, Mattfeld, Stark, & Stark, 2011). In this case, cultural differences would be reduced with age due to the inability to implement culturally-specific strategies that are cognitively demanding (Gutchess & Cho, 2024; Park et al., 1999). In other words, only age effects would be found in behavioral performance and in hippocampal engagement (e.g., Leal et al., 2017; Reagh et al. 2018; Yassa et al., 2011). The cultural differences identified for younger adults in their recruitment of LIFG in Leger et al. (2024a) would be reduced or eliminated with age. The current study will investigate joint effects of age and culture on both regions, as well as in the whole-brain using exploratory analyses. To our knowledge, this study is the first study to examine the joint effects of culture and age on memory using neural measures.

## Method

### Participants

Two hundred twenty-eight participants (60 American younger, 58 Taiwanese younger, 50 American older, and 60 Taiwanese older adults) completed the Mnemonic Similarity Task (MST) during MR scanning. Out of the 228, 21 participants were excluded from data analyses for the following reasons: excessive motion during the scan (please see the “data acquisition and preprocessing” section for details; *n* = 5), low performance during the MST (i.e., old vs. new discrimination was below chance or the number of trials of correct rejections for similar items was less than 5; *n* = 5), brain structural abnormalities (e.g., fissures in the frontal cortex; *n* = 4), a coil issue during the scan (*n* = 1), failure to meet inclusion guidelines (e.g., Americans with Asian ethnicity; *n* = 2), and procedural errors (i.e., incorrect files run or missing data files; *n* = 4). Thus, the final sample consisted of 55 American younger adults, 54 Taiwanese younger adults^1^, 43 American older adults^2^, and 55 Taiwanese older adults.

Based on a power analysis for a 2 (Age: younger vs. older) x 2 (Culture: American vs. Taiwanese) x 2 (Response to similar items: correct rejection vs. false alarm) repeated measures ANOVA using the G*Power (Faul et al., 2007), the required sample size for a medium effect size (f = 0.25), alpha = 0.05, and 1-β = 0.8 was at least 34 participants per group. We aimed to recruit larger sample sizes to detect stable and robust cultural effects.

Regarding culture, American and Taiwanese in the current study were defined as people who were born in the United States and Taiwan, respectively, and who had not lived in another country for more than 5 years. Americans were recruited from Brandeis University and the greater Boston area; Asian Americans were excluded due to the possibility of being exposed to elements of both cultures. Taiwanese were recruited from National Taiwan University and the surrounding Taipei area. Please see Table 1 for detailed demographic information of the sample.

**Table 1.**
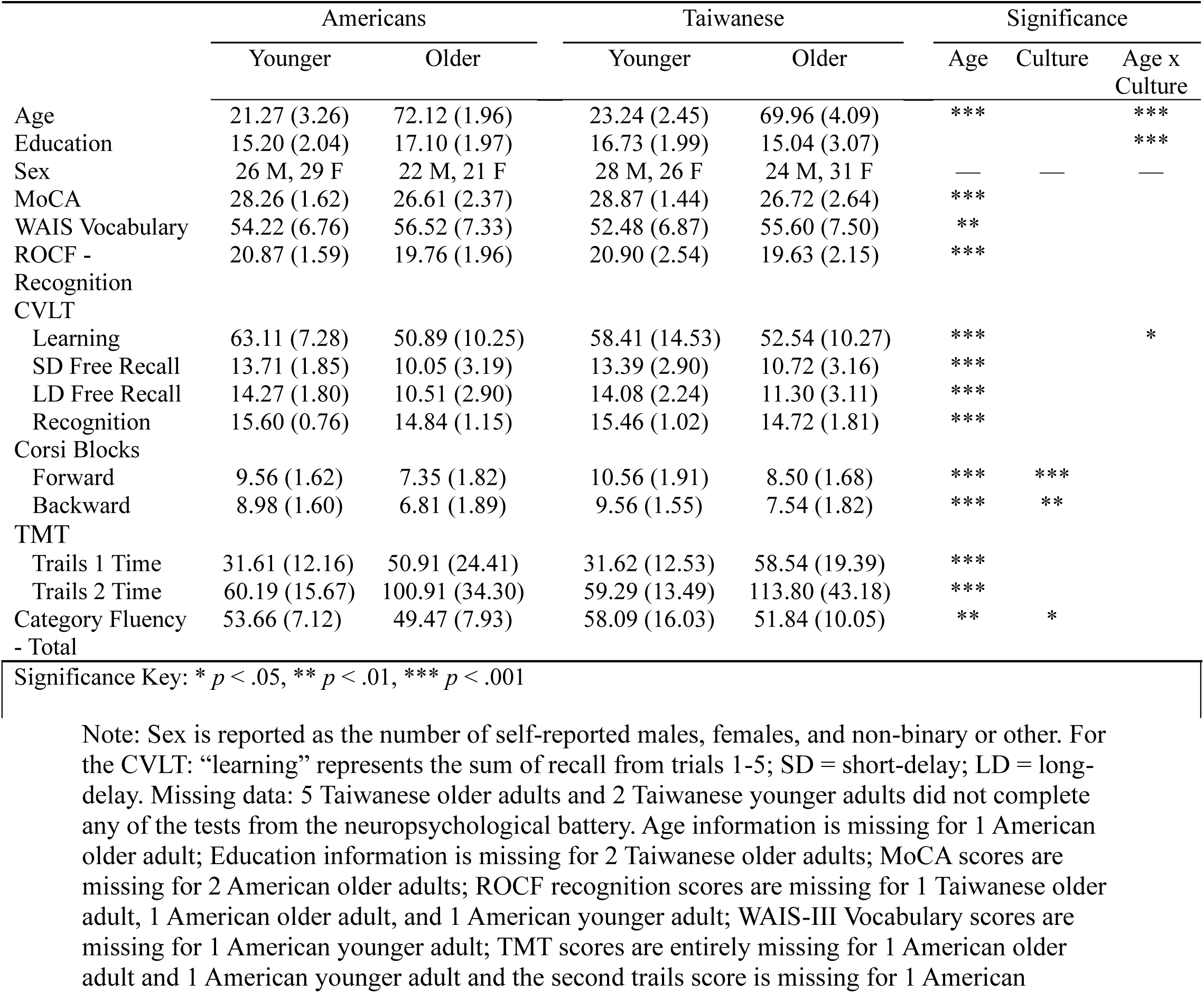
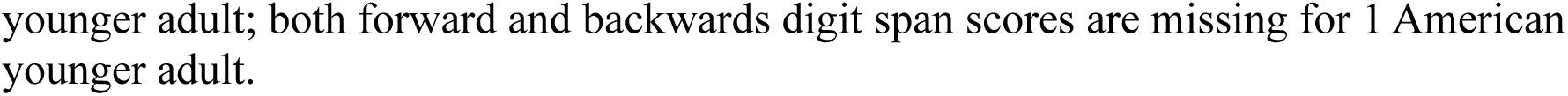
Means and Standard Deviations of Demographic Information and Neuropsychological Test Scores of Younger and Older American and Taiwanese Participants.

|  | Americans |  | Taiwanese |  | Significance |  |  |
| --- | --- | --- | --- | --- | --- | --- | --- |
|  | Younger | Older | Younger | Older | Age | Culture | Age x Culture |
| Age | 21.27 (3.26) | 72.12 (1.96) | 23.24 (2.45) | 69.96 (4.09) | *** |  | *** |
| Education | 15.20 (2.04) | 17.10 (1.97) | 16.73 (1.99) | 15.04 (3.07) |  |  | *** |
| Sex | 26 M, 29 F | 22 M, 21 F | 28 M, 26 F | 24 M, 31 F | — | — | — |
| MoCA | 28.26 (1.62) | 26.61 (2.37) | 28.87 (1.44) | 26.72 (2.64) | *** |  |  |
| WAIS Vocabulary | 54.22 (6.76) | 56.52 (7.33) | 52.48 (6.87) | 55.60 (7.50) | ** |  |  |
| ROCF - Recognition | 20.87 (1.59) | 19.76 (1.96) | 20.90 (2.54) | 19.63 (2.15) | *** |  |  |
| CVLT |  |  |  |  |  |  |  |
| Learning | 63.11 (7.28) | 50.89 (10.25) | 58.41 (14.53) | 52.54 (10.27) | *** |  | * |
| SD Free Recall | 13.71 (1.85) | 10.05 (3.19) | 13.39 (2.90) | 10.72 (3.16) | *** |  |  |
| LD Free Recall | 14.27 (1.80) | 10.51 (2.90) | 14.08 (2.24) | 11.30 (3.11) | *** |  |  |
| Recognition | 15.60 (0.76) | 14.84 (1.15) | 15.46 (1.02) | 14.72 (1.81) | *** |  |  |
| Corsi Blocks |  |  |  |  |  |  |  |
| Forward | 9.56 (1.62) | 7.35 (1.82) | 10.56 (1.91) | 8.50 (1.68) | *** | *** |  |
| Backward | 8.98 (1.60) | 6.81 (1.89) | 9.56 (1.55) | 7.54 (1.82) | *** | ** |  |
| TMT |  |  |  |  |  |  |  |
| Trails 1 Time | 31.61 (12.16) | 50.91 (24.41) | 31.62 (12.53) | 58.54 (19.39) | *** |  |  |
| Trails 2 Time | 60.19 (15.67) | 100.91 (34.30) | 59.29 (13.49) | 113.80 (43.18) | *** |  |  |
| Category Fluency - Total | 53.66 (7.12) | 49.47 (7.93) | 58.09 (16.03) | 51.84 (10.05) | ** | * |  |

Significance Key: \* $p < .05$ , \*\* $p < .01$ , \*\*\* $p < .001$
Note: Sex is reported as the number of self-reported males, females, and non-binary or other. For the CVLT: “learning” represents the sum of recall from trials 1-5; SD = short-delay; LD = long-delay. Missing data: 5 Taiwanese older adults and 2 Taiwanese younger adults did not complete any of the tests from the neuropsychological battery. Age information is missing for 1 American older adult; Education information is missing for 2 Taiwanese older adults; MoCA scores are missing for 2 American older adults; ROCF recognition scores are missing for 1 Taiwanese older adult, 1 American older adult, and 1 American younger adult; WAIS-III Vocabulary scores are missing for 1 American younger adult; TMT scores are entirely missing for 1 American older adult and 1 American younger adult and the second trails score is missing for 1 American
younger adult; both forward and backwards digit span scores are missing for 1 American younger adult.

**Table 2.** Means and standard deviations (SD) for response bias (c) across age and culture groups.

|  | Americans |  |  |  | Taiwanese |  |  |  |
| --- | --- | --- | --- | --- | --- | --- | --- | --- |
|  | Younger |  | Older |  | Younger |  | Older |  |
|  | Mean | SD | Mean | SD | Mean | SD | Mean | SD |
| Target-Foil | 0.20 | 0.31 | 0.22 | 0.37 | 0.21 | 0.31 | 0.18 | 0.45 |
| Target-Lure | -0.59 | 0.32 | -0.70 | 0.48 | -0.58 | 0.36 | -0.51 | 0.48 |

### Neuropsychological Tests

Participants completed a series of neuropsychological tests in a separate session prior to the scan. The neuropsychological battery included the Montreal Cognitive Assessment (MoCA; Nasreddine et al., 2005), the Rey–Osterrieth complex figure (ROCF; Meyers, 1994; Osterrieth, 1944), the Wechsler Adult Intelligence Scale (WAIS)-III Vocabulary (Wechsler, 1997), the California Verbal Learning Test Second Edition (CVLT-II, Delis et al., 2000), the Corsi block-tapping test (Corsi, 1972), Trail Making Test-Color (TMT; D’Elia et al., 1996), and category fluency test (Lezak, Howieson, Loring, Hannay, & Fischer, 2004; Hua, Chang, & Chen, 1997). Based on consultation with a neuropsychologist, tests were chosen to sample a number of cognitive domains (e.g., memory, executive function) and to minimize cultural bias, drawing on tests that have been used successfully across cultures. Please see Table 1 for scores on neuropsychological tests.

### Mnemonic Similarity Task (MST)

During scanning, participants completed the Mnemonic Similarity Task (MST, Kirwan, Jones, Miller, & Stark, 2007; Stark et al., 2015). The MST consisted of two phases (see Figure 1 for the structure of the task and example stimuli). First, in the encoding phase, participants saw 128 images of objects (e.g., balloons, flowers, frying pan) presented one at a time for 4 seconds. To ensure participants were attending to the stimuli, they decided whether each object belonged indoors or outdoors by pressing keys on the button box. Due to time constraints, structural brain images were acquired during the encoding phase. Next, participants completed a resting-state scan in which they viewed a fixation cross on the screen for ∼7 minutes^3^. After that, they completed the test phase of the MST as a surprise recognition test, and fMRI data were collected. In this phase, participants viewed 192 images of objects one at a time for 4 seconds followed by jittered fixation (between 800 ms to 12000 ms) and had to decide whether they had seen an object previously (i.e., “old”) or they had not (i.e., “new”) by pressing a button on the button box. Out of the 192 images, 64 images were the objects that participants had seen during the encoding phase (i.e., *Targets*), another 64 images were the objects similar to the objects that they had seen in the encoding phase (i.e., *Lures*), and the other 64 images were new objects (i.e., *Foils*). They were instructed to call lure objects “new.” There were 4 runs in the test phase, and each run took about 6 minutes and 31 seconds.

**Figure 1.**
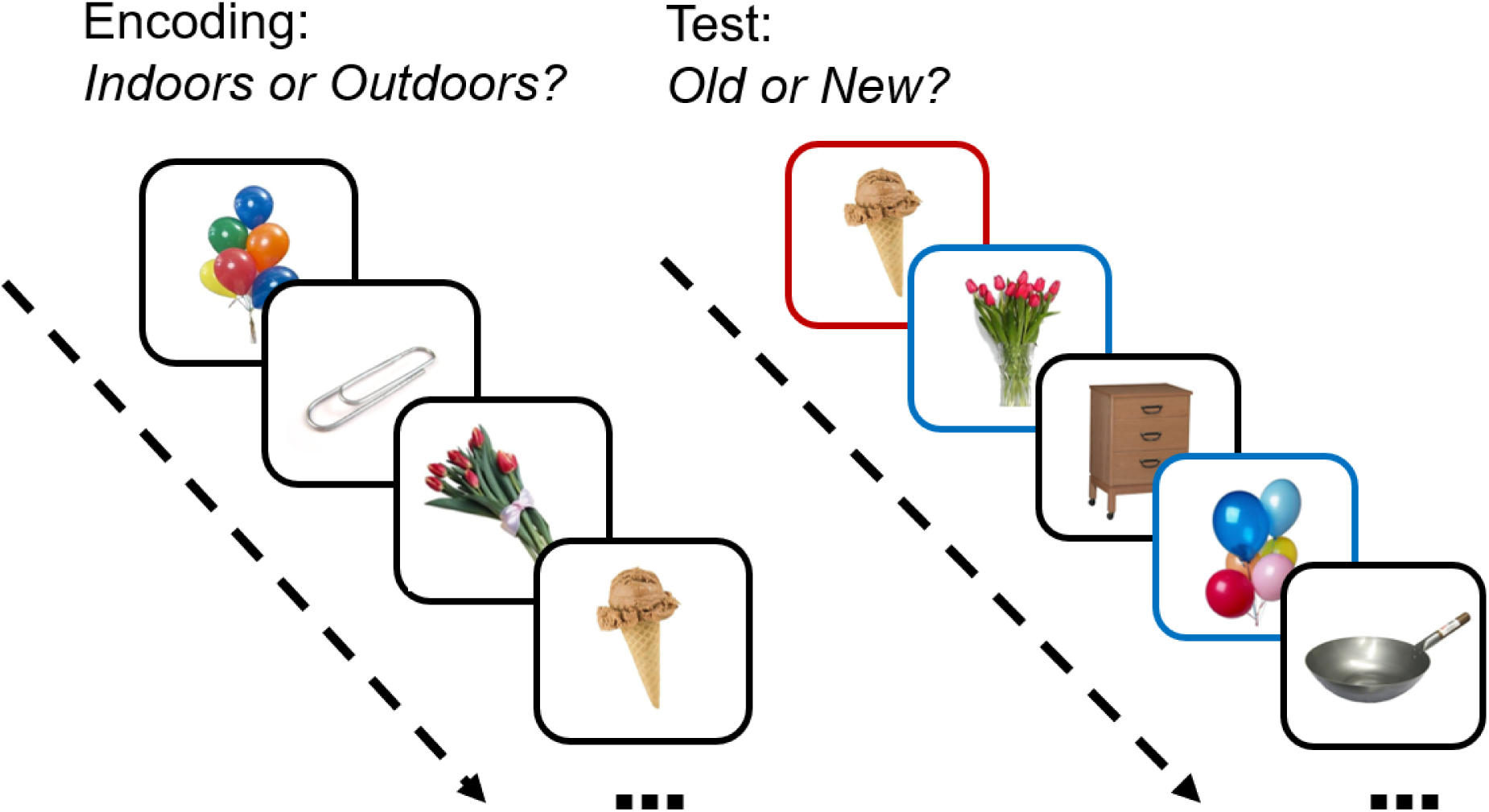
The task structure and example stimuli for the Mnemonic Similarity Task (MST). In this example, the ice cream cone is an example of a target (denoted with a red frame at test, for illustration purposes in this figure), the flowers and balloons are examples of lures (denoted with blue frames), and the dresser and frying pan are the examples of foils (denoted with black frames).

For behavioral measures, we used d’ and c scores. The d’ scores indicate memory sensitivity to discriminate old from new items, calculated as the z score of false-alarms – z score of hits. We calculated Target-Foil d’ (i.e., z score of “old”|foils – z score of “old”|targets) and Target-Lure d’ (i.e., z score of “old”|Lures – z score of “old”|targets), measuring one’s sensitivity to discriminate targets and foils and discriminate targets and lures, respectively. Therefore, Target-Lure d’ is the most relevant measure to pattern separation. Response bias c scores indicate one’s tendency to respond “old” or “new,” calculated by averaging z scores of hits and those of false-alarms and then multiplying the averaged z scores by -1. If the c scores are positive values, it means individuals have greater tendency to respond “new,” whereas if the c scores are negative values, it means individuals have greater tendency to respond “old.” Because scores cannot be calculated when values are at ceiling or floor, rates of 1 were adjusted to (N_trials_ -1)/ N_trial_ and rates of 0 were adjusted to 1/N_trials_.

### Neuroimaging Data

#### Data acquisition and preprocessing

In both the U.S. and Taiwan, 3T Siemens MAGNETOM Prisma whole-body MRI systems (Siemens Medical Solutions, Erlangen, Germany) were used. In the U.S., neuroimaging data was collected at Center for Brain Science at Harvard University, Cambridge, MA, and in Taiwan, the data was collected at the Imaging Center for Integrated Body, Mind, and Culture Research at National Taiwan University, Taipei.

We first conducted calibration scans with the same participants tested across sites to ensure no meaningful differences in global signal occurred across the scanners (Chen et al., 2020). To obtain images, we used a 64-channel head coil and simultaneous multi-slice scanning (Moeller et al., 2010, Xu et al., 2013), which enabled us to acquire 2.3 mm thick slices with whole-brain coverage using an echo-planar image sequence (with TE = 25ms, TR = 800 ms, FOV = 220 mm, and flip angle = 60°). For high resolution T1-weighted images, a multi-echo MPRAGE sequence (van der Kouwe et al., 2008) was used to acquire 176 1.0 x 1.0 x 1.0 mm slices (with short TE = 1.69 ms, long TE = 7.27 ms, TR = 2,530.0 ms, FOV = 256 X 256 mm, and FA = 7°) were acquired. For preprocessing, we used fMRIPrep 20.0.6 (Esteban et al., 2019); please see Leger et al. (2024a) for more details on data acquisition and preprocessing.

The current analyses used the following criteria to define excessive motion: participants whose average of framewise displacement across all runs was greater than 0.5 or whose averaged number of images flagged as motion outliers during preprocessing pipeline across all runs was more than 30 % of the images contained in a single run (i.e., 139) were excluded. The criteria were set independently from those used in Leger et al. (2024a) to be appropriate for samples of older adults. After preprocessing steps, Diffeomorphic Anatomical Registration Through Exponentiated Lie algebra (DARTEL) (Ashburner, 2007) was used to create a template from the final sample of participants in order to have a culture- and age-fair brain template for normalization to MNI space.

#### General linear model

We used SPM12 (Wellcome Department of Cognition Neurology, London, UK) for analyses. At the first-level, five behavioral regressors were constructed: hits for targets (“old”|Targets), correct rejections for lures (“new”|Lures), false alarms to lures (“old”|Lures), correct rejections for foils (“new”|Foils), and a regressor of no interest collapsing misses for targets (“new”|Targets) and false alarms to foils (“old”|Foils), due to small numbers of trials in these conditions. Each trial was modeled using a delta function (i.e., stimulus onset; duration = 0) and convolved with the hemodynamic response function. Also, six motion vectors (i.e., x, y, z, pitch, roll, yaw) and the five largest anatomical component-based noise correction (aCompCor) components from preprocessing steps were included as regressors.

To assess neural activity related to pattern separation, contrasts between correct rejections of lures and false alarms to lures were created. We next ran a 2 (age: younger vs. older) x 2 (culture: American vs. Taiwanese) full factorial model in SPM12. To compare neural activity to discrimination of old from new stimuli, the same analytic approach was applied using the contrasts between hits for targets and correct rejections for foils.

We adopted a region of interest (ROI) approach to target regions for which we had the strongest a priori predictions and to aid in interpreting comparisons across the four groups.

Because regions of the hippocampus are involved in pattern separation (e.g., Leal & Yassa, 2018; Yassa & Stark, 2011), we created a bilateral hippocampus mask using the automatic anatomical labeling (AAL) atlas from the WFU PickAtlas Tool. In addition to examining the hippocampus, we examined the joint influences of age and culture on the response of left inferior frontal gyrus (LIFG) as a second region of interest. This region was selected because cultures differed in the recruitment of the region in the previous comparison of old vs. new memory in younger adults (Leger et al., 2024a). We probed the LIFG to examine whether the pattern of cultural differences seen in younger adults extended to older adults or whether aging mitigated these cultural differences. To do so, we created a LIFG mask based on the region that emerged in the comparison of younger adults across cultures. That is, we re-ran the contrast that yielded cultural differences in the paper comparing younger adults (Leger et al., 2024a) in our sample of younger adults (*see* footnote 1): old > new discrimination (hits for targets vs. correct rejections for foils) in Younger Taiwanese > Younger Americans. Using a threshold of p < .001 and k = 116, calculated based on AFNI 3dClustSim algorithm (Cox & Hyde, 1997; Cox et al., 2017) for old vs. new discrimination to achieve a corrected p < .05, a cluster including the LIFG was the only region to emerge (k = 558). Thus, we generated a functional ROI mask from this region, which includes triangular and orbital IFG as well as left frontal operculum [peak voxel: -41, 25, 0], Brodmann’s areas: BA 47, 45, 38). Because this mask was generated using the younger adult data from this sample, the focus will be on comparing older adults across cultures; data from younger adults will be included for comparison with the older adults.

To conduct ROI analyses, we extracted betas from each of the two ROIs for each participant, and then ran 2 (age: younger vs. older; between-subject) x 2 (culture: American vs. Taiwanese; between-subject) x 2 (condition; the analysis of pattern separation compares correct rejections vs. false alarms for lures and the analysis of old vs. new discrimination compares hits for targets vs. correct rejections of lures). Both comparisons of memory conditions were conducted in both ROIs. ANOVAs were run using IBM SPSS Statistics (Version 29).

In addition to the ROI comparisons, we also conducted exploratory whole-brain analyses. Given the absence of fMRI data on cross-cultural comparisons of memory, particularly including samples of older adults, this approach allows for the identification of additional regions that are engaged differently across culture and age groups. We used the criteria of p(unc.) < .005 for voxel thresholds and k = 299 for pattern separation and p(unc.) < .005 for voxel thresholds and k = 318 for old vs. new discrimination to achieve a whole-brain cluster-wise family-wise error of p(FWE) < .05. These are calculated based on AFNI 3dClustSim algorithm. Given the absence of significant whole-brain effects using the above cluster sizes, a more lenient threshold of *p* = .001 (uncorrected) at the voxel level and k = 10 was also used.

## Results

### Behavioral Tasks

#### Demographics and Neuropsychological Tasks

Table 1 reports scores for each group, and the results of 2 x 2 ANOVAs comparing age and culture groups. Aside from the WAIS Vocabulary task, for which older adults had higher scores than younger adults, younger adults had higher or faster scores than older adults. Although the Taiwanese younger adults were slightly older with more years of education compared to the American younger adults and the Taiwanese older adults slightly younger with fewer years of education compared to the American older adults, the samples were overall well-matched across cultures on cognitive ability.

Taiwanese had higher scores than Americans on the Corsi blocks and Category Fluency Task, there was a larger age differences CVLT learning scores for Americans compared to Taiwanese, but there were no cultural differences in tests of cognitive orientation (MoCA), vocabulary (WAIS Vocabulary), long-term memory (ROCF and CVLT), or executive function (TMT).

**MST.** A 2 (age: younger vs. older; between-subject) x 2 (culture: American vs. Taiwanese; between-subject) x 2 (memory condition: Target-Foil vs. Target-Lure; within-subject) analysis of variance (ANOVA) was conducted. The dependent variables were memory discrimination (d’) and response bias (c).

***Memory discrimination (d’).*** There was a significant main effect of age, *F*(1, 203) = 48.20, *p* < 0.001, 𝜂_p_^2^ = 0.19, such that younger adults had higher levels of memory performance than older adults. The significant main effect of culture, *F*(1, 203) = 20.04, *p* < 0.001, 𝜂_p_^2^ = 0.09, occurred due to Americans’ overall higher level of memory performance compared to Taiwanese. There was a main effect of condition such that memory performance in the Target-Foil condition was higher than in the Target-Lure condition, *F*(1, 203) = 1575.04, *p* < 0.001, 𝜂_p_^2^ = 0.89. In terms of interactions, there was a significant interaction of Culture x Memory Condition, *F*(1,203) = 9.46, *p* < 0.005, 𝜂_p_^2^ = 0.05, and critically, a significant interaction of Age x Culture x Memory Condition, *F*(1,203) = 7.49, *p* < 0.01, 𝜂_p_^2^ = 0.04. No other interactions approached significance, *p*s > 0.18. Results are displayed in Figure 2.

**Figure 2.**
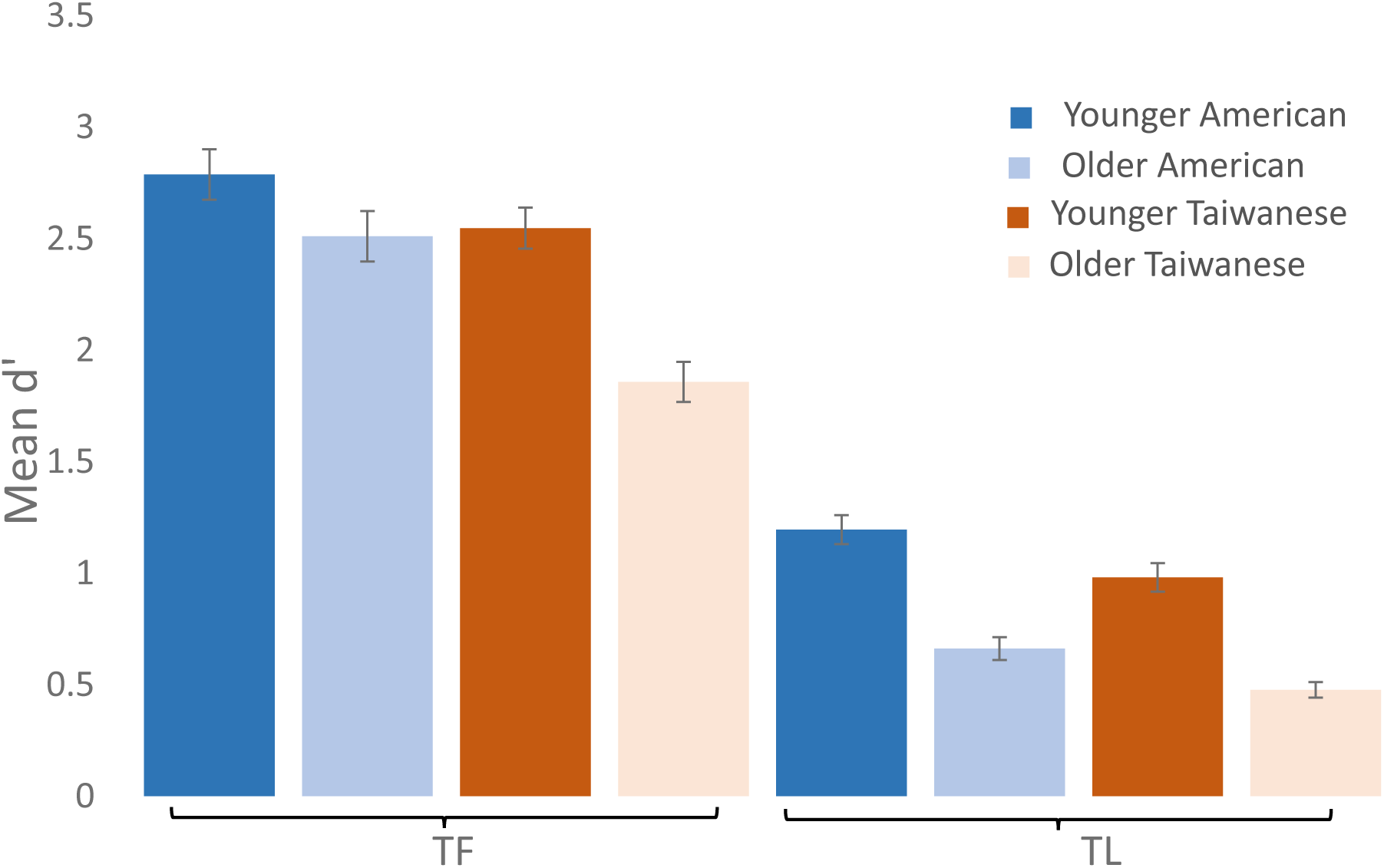
Memory discrimination (d’) for the American and Taiwanese younger and older adults for conditions TF (Target-Foil) and TL (Target-Lure) on the Mnemonic Similarity Task (MST). Error bars represent standard errors.

To further understand the nature of the 2×2×2 interaction, we conducted two separate 2 (age: younger vs. older; between-subject) x 2 (culture: American vs. Taiwanese; between-subject) ANOVAs in order to compare the influence of these factors on each memory condition. For the Target-Foil ANOVA, there were main effects of age, *F*(1, 203) = 22.28, *p* < 0.001, 𝜂_p_^2^ = 0.10, and culture, *F*(1, 203) = 19.02, *p* < 0.001, 𝜂_p_^2^= 0.09. The interaction of age x culture was significant, *F*(1, 203) = 4.05, *p* < 0.05, 𝜂_p_^2^ = 0.02. To further understand the nature of the interaction, we broke it down with two independent samples t-tests comparing the age groups within each culture. For the Americans, there was no significant difference between the performance of the younger and older adults, *t*(96) = 1.71, *p* = 0.09. In contrast, the younger Taiwanese had higher levels of memory performance than the older Taiwanese, *t*(107)= 5.36, *p* < 0.001.

For the Target-Lure ANOVA, both main effects of culture, *F*(1, 203) = 12.56, *p* < 0.001, 𝜂_p_^2^ = 0.06, and age, *F*(1, 203) = 85.02, *p* < 0.001, 𝜂_p_^2^ = 0.30, were significant, but the interaction

*Memory response bias (c).* There was a significant main effect of condition such that participants were less likely to report seeing “old” items in the Target-Foil condition than in the Target-Lure condition, *F*(1,203) = 1575.04, *p* < 0.001, 𝜂_p_^2^ = 0.89. In terms of interactions, there was a significant interaction of culture x memory condition, *F*(1,203) = 9.46, *p* < 0.005, 𝜂_p_^2^ = 0.05, and critically, a significant interaction of age x culture x memory condition, *F*(1,203) = 7.49, *p* < 0.01, 𝜂_p_^2^ = 0.04. No other main effects or interactions approached significance, *p*s > 0.41.

To further understand the nature of the 2×2×2 interaction, we separately analyzed each age group by conducting two 2 (culture: American vs. Taiwanese; between-subject) x 2 (memory condition: Target-Foil vs. Target-Lure; within-subject) ANOVAs. For the younger adults, there was a significant main effect of memory condition, *F*(1, 107) = 793.47, *p* < 0.001, 𝜂_p_^2^ = 0.88, but neither the main effect of culture nor the interaction of culture x memory condition approached significance, *ps* > 0.80. The older adults also exhibited a significant main effect of memory condition, *F*(1, 96) = 788.34, *p* < 0.001, 𝜂_p_^2^ = 0.89, but the main effect of culture did not approach significance, *p* > .40. Critically, there was a significant interaction of culture x memory condition, *F*(1, 96) = 16.55, *p* < 0.001, 𝜂_p_^2^ = 0.15. Follow-up independent samples t-tests revealed that the American older adults’ Target-Lure c values tended to be lower than their Taiwanese counterparts (i.e., American older adults displayed greater tendency to respond “old,” having a more liberal bias, compared to Taiwanese older adults), *t*(96) = 1.94, *p* <.06, although this effect did not reach traditional levels of significance. In contrast, there was no evidence of cultural difference in older adults for the c values in the Target-Foil c condition, *p* = .60.

### Functional MRI

To examine the joint effects of age and culture, we conducted a priori ROI analyses to investigate neural activity related to pattern separation as well as old/new discrimination. In addition, we conducted whole-brain exploratory analyses.

## ROI analyses

### Pattern separation (correct rejections vs. false alarms for lure items)

Two ROIs (i.e., bilateral hippocampus and LIFG) were used for ROI analyses (*see* Figure 3). For the bilateral hippocampus ROI, none of the effects approached significance (*p*s > .10). For the LIFG ROI, results are shown in Figure 4. Most importantly, there was no significant interaction of age x culture x condition, *F*(1, 203) = 0, *p* = 1.00. However, we observed a significant interaction of culture x condition, *F*(1, 203) = 6.74, *p* = .01, 𝜂_p_^2^ = 0.03. Post-hoc analysis with a Bonferroni adjustment (estimated marginal means) showed that Americans had significantly higher activation in this region for correct rejections compared to false alarms (*p* = .004), whereas brain activity between correct rejections and false alarms did not differ for Taiwanese (*p* = .49). In addition, there was a significant interaction of age x condition, *F*(1, 203) = 5.34, *p* = .022, 𝜂_p_^2^ = 0.03. Specifically, younger adults showed significantly greater brain activity for correct rejections compared to false alarms (*p* = .004), but the conditions did not significantly differ for older adults (*p* = .65). The main effect of age was also significant, *F*(1, 203) = 15.32, *p* < .001, 𝜂_p_^2^ = 0.07, such that older adults demonstrated significantly more activity overall compared to younger adults. No other effects were significant, *p*s > .06.

**Figure 3.**
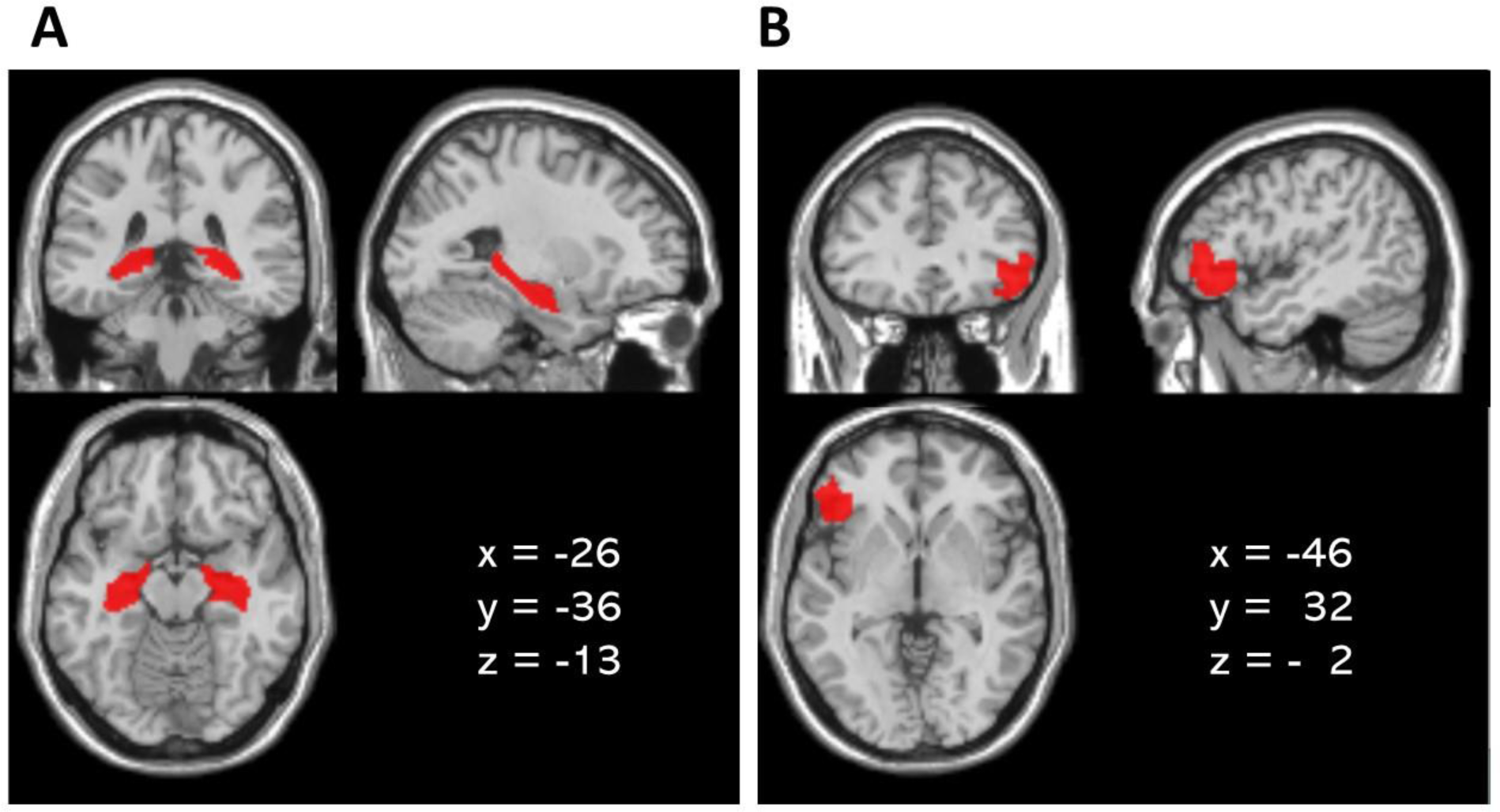
The illustration of ROIs: (A) the bilateral hippocampus structural ROI and (B) the LIFG ROI defined based on younger adults’ functional data.

**Figure 4.**
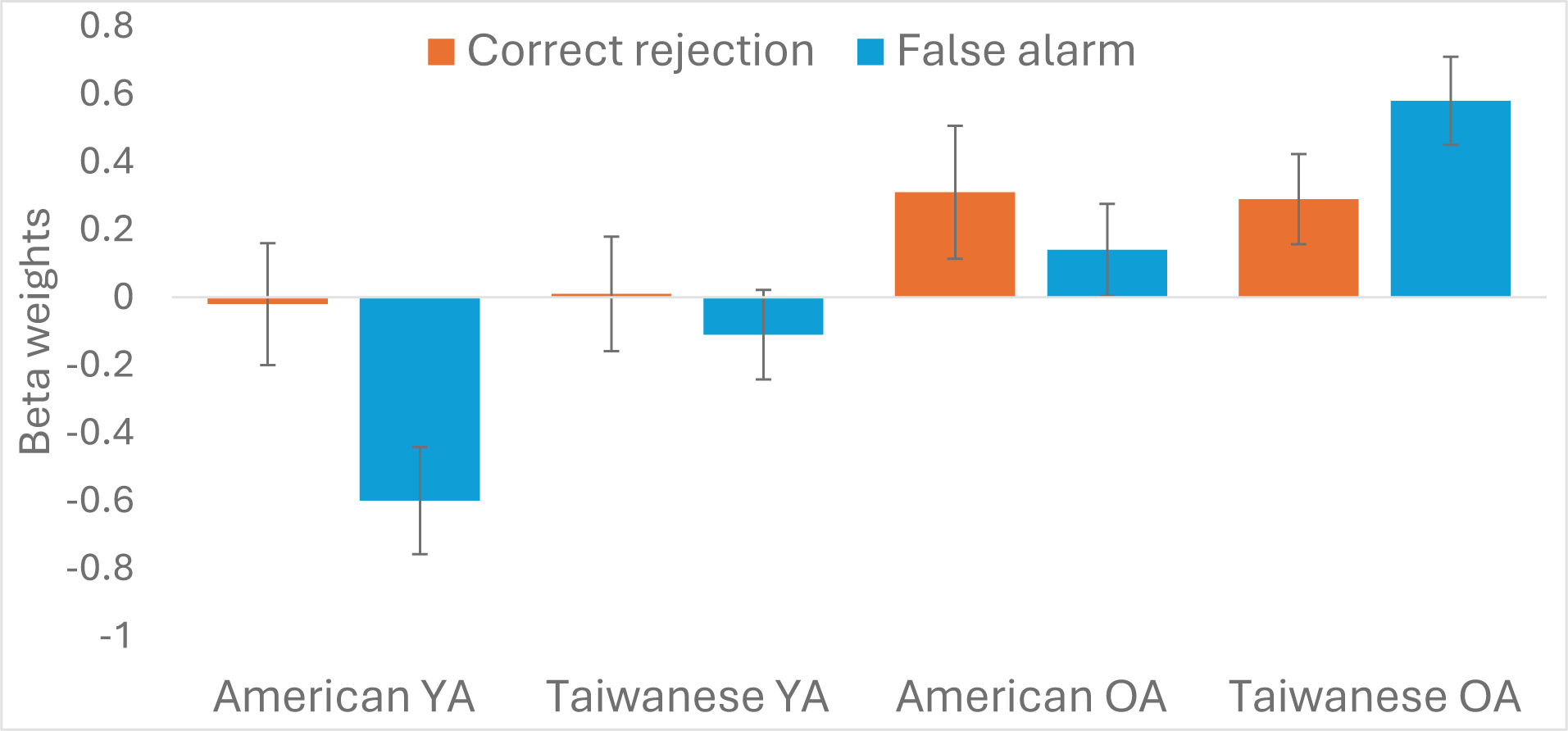
Beta weights from the LIFG ROI mask for pattern separation (i.e., correct rejections vs. false alarms for lures) for each group. Error bars represent standard errors.

### Old vs. New (hits for targets vs. correct rejections for foils)

For the bilateral hippocampal ROI, results are shown in Figure 5. Most importantly, there was no significant interaction of age x culture x condition, *F*(1, 203) = 0.14, *p* = .71. However, we found a significant interaction of culture x condition, *F*(1, 203) = 6.70, *p* = .01, 𝜂_p_^2^ = 0.03, such that Americans displayed greater activity for correct rejections of foils compared to hits to targets (*p* = .008), whereas such differences were not found in Taiwanese (*p* = .34). This effect emerged in the previous analysis of younger adult data across cultures (Leger et al., 2024a); these analyses demonstrate that the pattern extends across younger and older adults. None of the other effects were significant, *p*s > .05.

**Figure 5.**
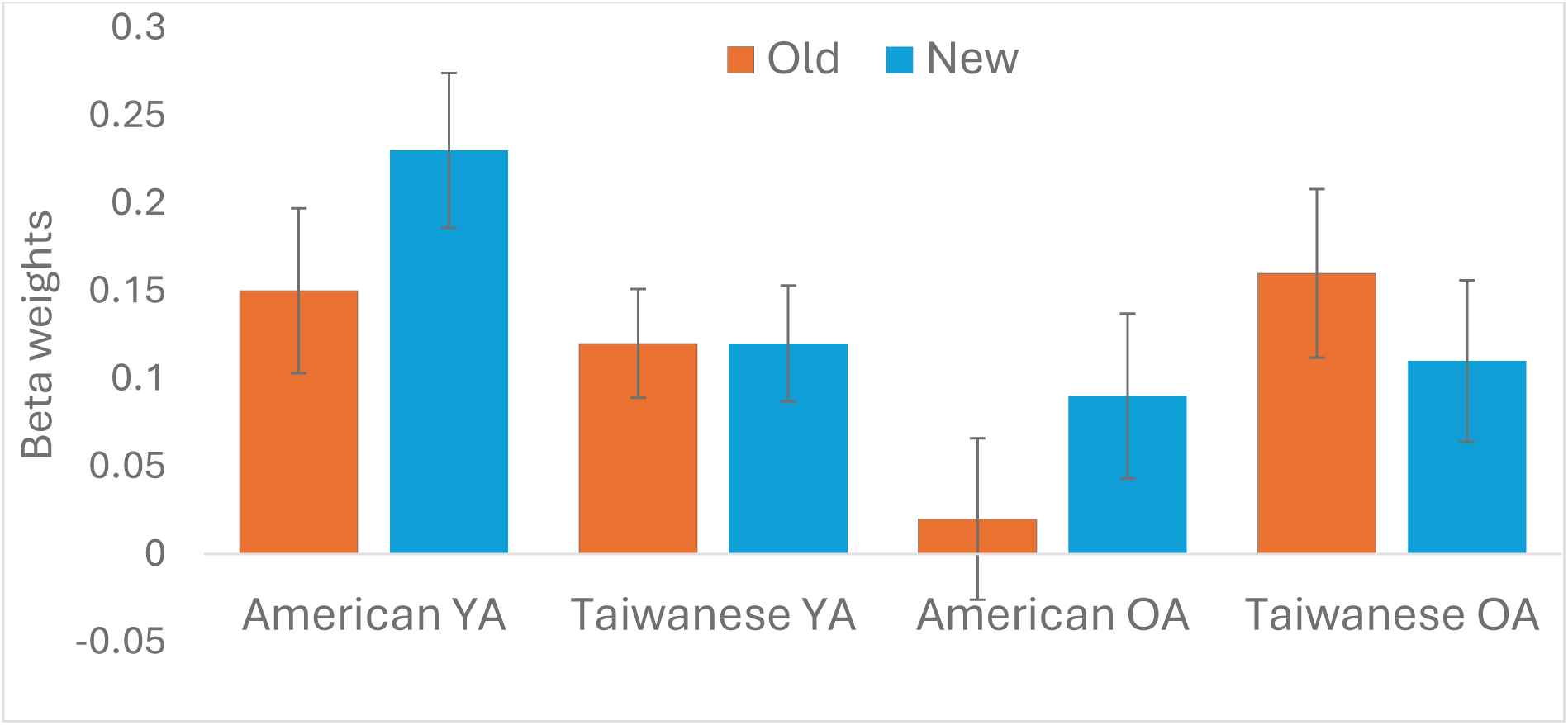
Beta weights from the bilateral hippocampus ROI mask for old vs. new discrimination (i.e., hits for targets vs. correct rejections for foils) for each group. Error bars represent standard errors.

For the LIFG ROI, results are shown in Figure 6. Critically, there was a significant age x culture x condition interaction, *F*(1, 203) = 7.87, *p* = .006, 𝜂_p_^2^ = 0.04. Specifically, American younger adults displayed greater activation for correct rejections of foils than hits for targets (*p* < .001), whereas the opposite pattern was found in Taiwanese younger adults (*p* = .002). However, in older adults from both countries, the activations were not significantly different between hits for targets and correct rejections for foils (*p*s > .06). Extending the previous research based on younger adults (Leger et al., 2024a), the results suggest that the cross-cultural differences present in younger adults do not extend to older adults. A significant interaction of culture x condition was also observed, *F*(1, 203) = 20.33, *p* < .001, 𝜂_p_^2^ = 0.09, such that Americans had higher levels of activation for correct rejections of foils compared to hits for targets (*p* = .004), whereas Taiwanese showed the opposite pattern (*p* < .001). The main effect of age was also significant, *F*(1, 203) = 38.48, *p* < .001, 𝜂_p_^2^ = 0.16. Specifically, older adults displayed higher levels of overall activations than younger adults. None of the other effects were significant, *p*s > .11.

**Figure 6.**
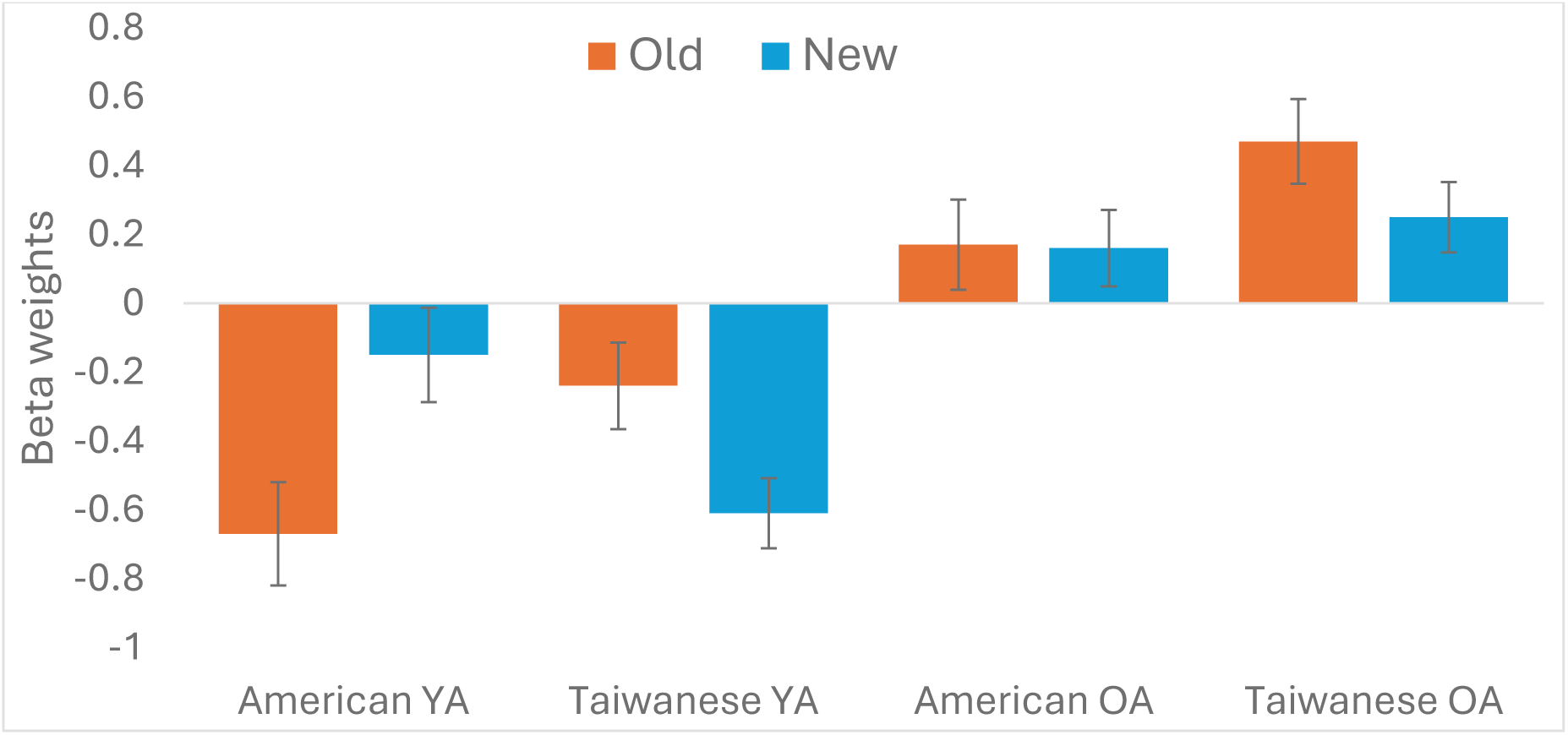
Beta weights from the LIFG ROI mask for old vs. new discrimination (i.e., hits for targets vs. correct rejections for foils) for each group. Error bars represent standard errors.

## Exploratory Whole-Brain Analyses

### Pattern Separation

For the exploratory whole-brain analysis, no significant clusters emerged for the interaction of age by culture at a corrected *p* <.05. At a more lenient uncorrected threshold, using *p* = .001 and k = 10, some clusters showed an age by culture interaction. See Table 3 for details.

**Table 3.**
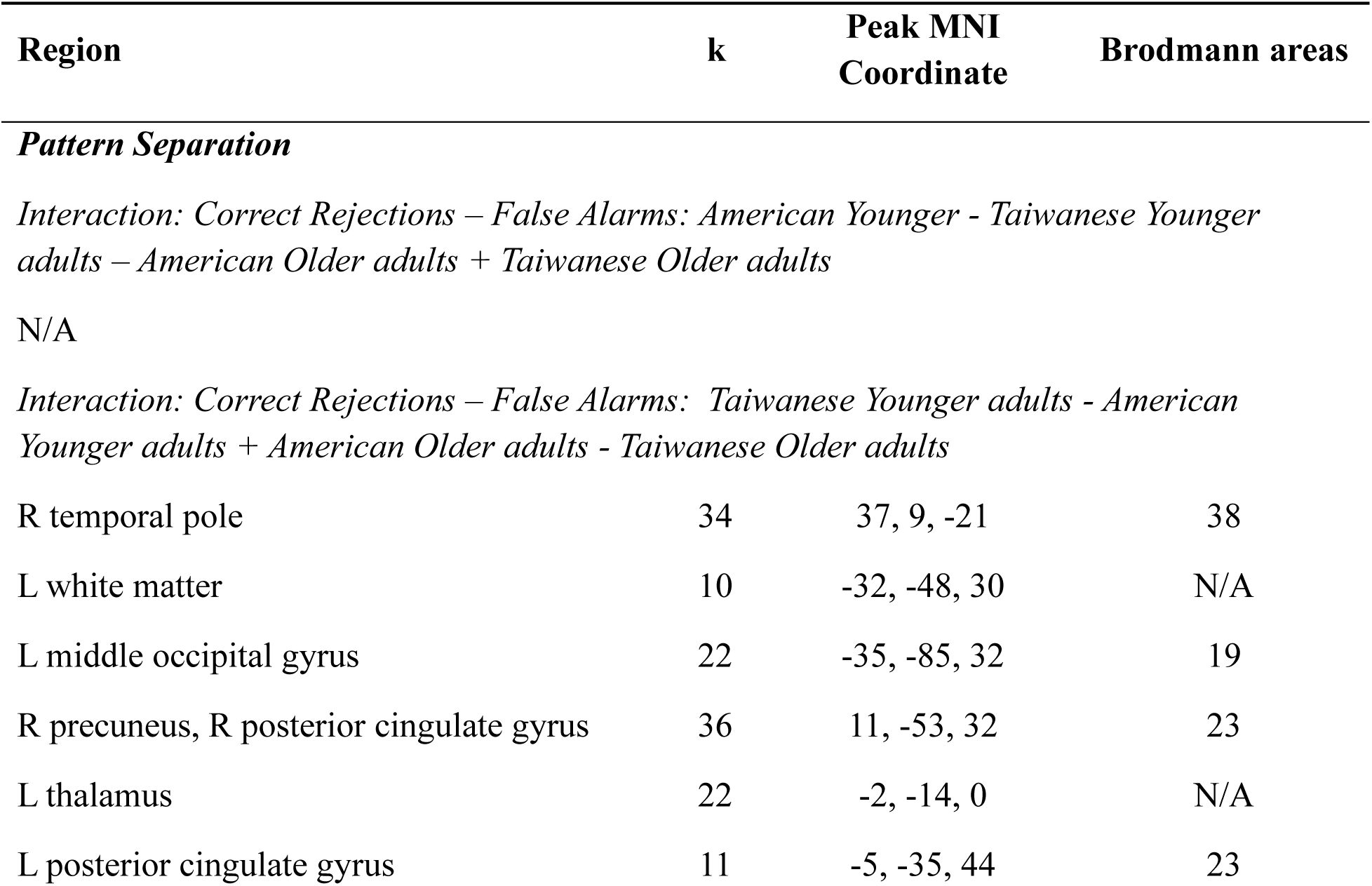
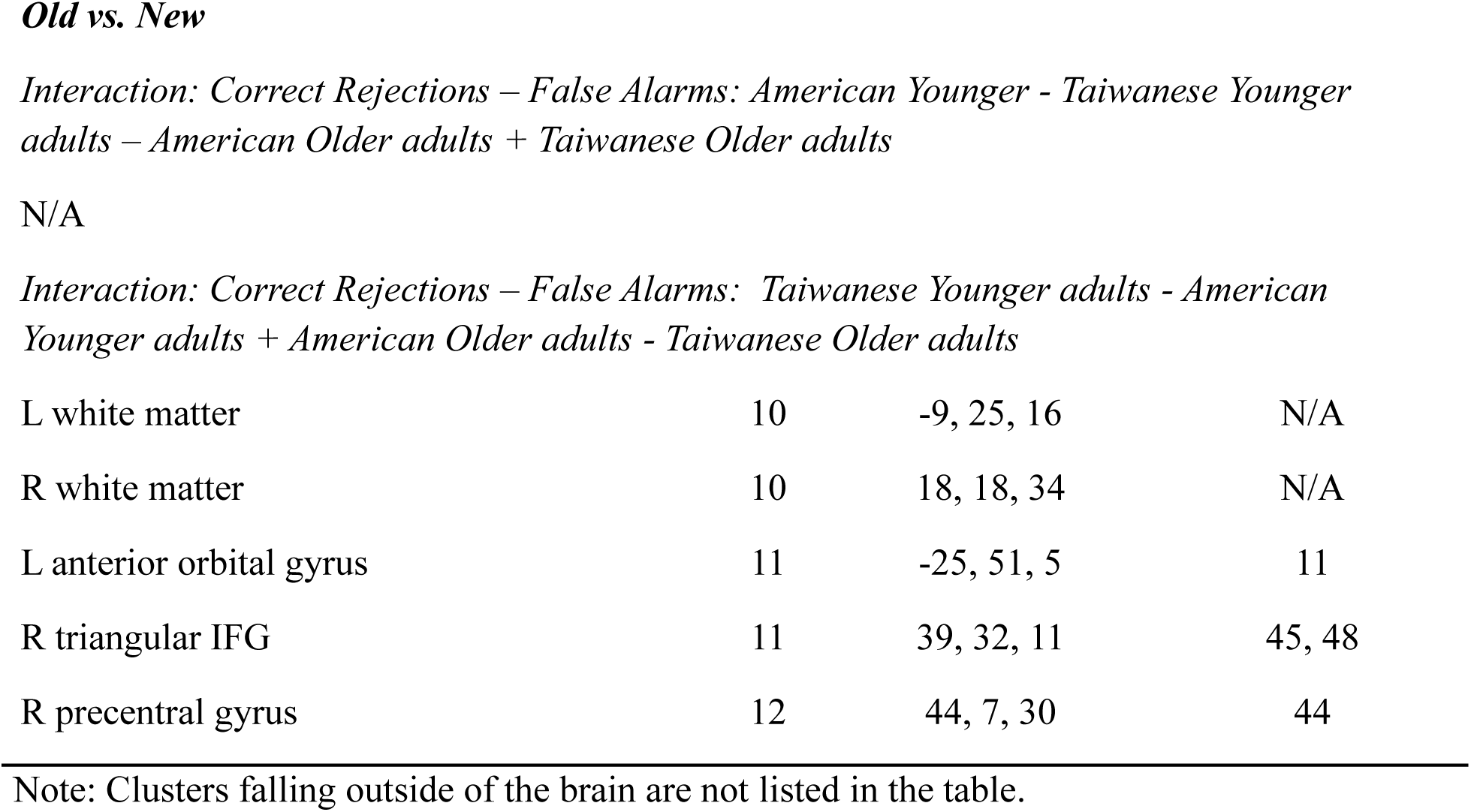
Regions showing an Age x Culture interaction in the exploratory whole-brain analysis for pattern separation (correct rejections vs. false alarms to lures) and old vs. new (hits for targets vs. correct rejections of foils) when using a threshold of p = .001 and k = 10. (No regions emerged as significant in the analyses at a corrected p<.05)

## Discussion

The current study examined joint effects of age and culture on memory using behavioral and neural measures. Extending a previous study that investigated cross-cultural differences in memory using the Mnemonic Similarity Task (Stark et al., 2019) to compare American and Taiwanese younger adults (Leger et al., 2024a), the present study added American and Taiwanese older adults. The present study makes five main contributions to the understanding of the combined effects of age and culture on memory.

First, we hypothesized that pattern separation, based on the comparison of correct rejections and false alarms to lures, would serve as a potential mechanism to account for cross-cultural differences in memory specificity. Behaviorally, Americans had higher levels of memory performance (d’) than Taiwanese on measures associated with pattern separation. This notably differs from our prior publication of the younger adult data (Leger et al., 2024a), which failed to detect cultural differences due to the inclusion of an additional (lure-foil) condition that prevented us from detecting this effect^4^. However, despite finding evidence for behavioral differences across cultures, we do not have strong support that cultural differences are specific to pattern separation. This is because neurally there were no effects of culture on hippocampal activity, which is the region most strongly associated with pattern separation (Leal & Yassa 2018; Yassa & Stark, 2011). The lack of cultural differences in the hippocampus was true whether collapsing across age groups or testing for interactions with age. Moreover, the results can be considered in light of the typical effects of age, regardless of culture. Older adults performed worse than younger adults on behavioral measures of pattern separation, consistent with the previous literature (Leal et al., 2017; Reagh et al. 2018; Stark et al., 2010, 2013, 2015; Yassa et al., 2011). However, our study failed to find age differences in hippocampal activity. We speculate that the lack of hippocampal findings reflects the need to adopt a more fine-grained approach to distinguish subfields of the hippocampus, rather than our blunter ROI approach.

Pattern separation analyses revealed a second effect. Cultures differed in LIFG activity such that Americans – collapsing across age groups - engaged the regions more for correct rejections compared to false alarms of lures whereas activity in the Taiwanese did not differ across the conditions. This finding extends the cultural difference in LIFG engagement identified in young adults during old vs. new judgments (Leger et al., 2024a) to pattern separation comparisons, identifying an overall cultural difference. However, it may be the case that we do not have the power to detect an interaction of age and culture on the engagement of these regions across conditions. Visually inspecting the activity across conditions (Figure 4), there is a tendency for the pattern to reverse for older adults – with activity for false alarms higher than for correct rejections for Taiwanese whereas there is the opposite trend for Americans. This suggests that older adults may drive the overall finding of cultural differences. Moreover, the interaction of age and condition indicated that younger adults engage this region for correct rejections more than false alarms, whereas the conditions do not differ for older adults. Thus, how age affects cultural differences in activity related to pattern separation in LIFG is an open question, though activity in this region consistently suggests Americans evidence more activity for correct rejections than false alarms. This could indicate Americans’ greater attention to novelty, as we will discuss for the following sections.

Comparisons of old vs. new items yields more straightforward extensions of prior research. Behaviorally discriminating old vs. new objects supported higher performance in Americans than Taiwanese, consistent with some previous findings with younger adults (Leger & Gutchess, 2021; Leger et al., 2024a), as well as higher levels of performance for younger compared to older adults (Fraundorf et al., 2019). Interestingly, the effects of age on old/new discrimination in memory were more pronounced for Taiwanese than Americans, which is the third major contribution of this study. This finding indicates differential vulnerability across cultures to age-related impairments in remembering visually detailed objects. Effects of age are pervasive, occurring for both cultural groups. However, it may be the case that Americans’ relatively greater emphasis on details in memory (Leger & Gutchess, 2021; Leger et al., 2024a; Millar et al., 2013) and analytic processing (Nisbett et al., 2001) in young adulthood could mitigate, to some extent, effects of age. This could suggest that a lifetime of experience with culturally-specific practices (Gutchess & Cho, 2024) or cognitive strategies could be an effective way to reduce some age-related detriments in memory (Craik, 2000; Salthouse, 2009; Zack, Hasher & Li, 2000).

Fourth, previous findings of cultural differences in hippocampal activity for correct new vs. old judgments (Leger et al., 2024a) extended to older adults. Collapsing across age groups, Americans showed higher levels of activation in the hippocampus for correct memory for new vs. old judgments, whereas activity in the Taiwanese did not differ across judgments. Results suggest that Americans may recruit the hippocampus more in response to novel than old information (Fredes & Shigemoto, 2021; Kumaran & Maguire, 2009) than Taiwanese do, even into older adulthood. We (Leger et al., 2024a) interpreted this cultural difference in younger adults as potentially reflecting differences in memory states, such as orienting to novelty (encoding) or familiarity (retrieval) (Long & Kuhl, 2021). This could mean that in response to novel items, Americans attempt pattern separation whereas Taiwanese attempt pattern completion; the current results suggest these tendencies could extend to older adulthood. However, direct tests using tasks appropriate to assess novelty orientation are needed to substantiate this interpretation.

Lastly, cultural differences in the activation of LIFG for correct old vs. new judgments in younger adults are eliminated with age. Specifically, when correctly retrieving memories, younger Americans engaged the LIFG for new items more than old ones, whereas younger Taiwanese engaged the LIFG more for old than new items (Leger et al., 2024a). With age, activity for old vs. new judgments did not differ across cultures. We had speculated that the pattern in younger adults could reflect that younger Americans might encode more detailed representations of targets, and thus experience less interference at retrieval when discerning old from new information (Leger et al., 2024a). In contrast, if younger Taiwanese encoded less detailed information, they could experience more interference in discerning old from new memories at retrieval. This interference could induce higher levels of activation in the LIFG for old vs. new judgments, reflecting the role of LIFG in cognitive control processes (Badre & D’Esposito, 2007; Badre & Wagner, 2005; Thompson-Schill, D’Esposito, Aguirre, & Farah, 1997). The absence of cultural differences in LIFG activity in older adults, particularly as activity was equivalent for old and new items in Americans, could reflect reductions in cognitive control with age (Manard, Carabin, Jaspar, & Collette, 2014; Salthouse, Atkinson, & Berish, 2003). Because this interpretation is based on inferences about the role of the region in cognitive control (Badre & D’Esposito, 2007; Badre & Wagner, 2005; Thompson-Schill, D’Esposito, Aguirre, & Farah, 1997), it is necessary to design a study to directly test this explanation, as well as compare encoding and retrieval processes within participants in order to assess trade-offs across cultures in these stages of memory. Nevertheless, the results indicate that the effects of age on LIFG activity may surpass cultural effects in that cultural differences present in younger adults are eliminated in older adults.

The behavioral and neural findings illustrate the mixed results of how age and culture interact in memory. In future research, it would be promising to further investigate how the availability of cognitive resources contribute to the relationship between age and culture. Existing frameworks propose that for tasks that demand fewer cognitive resources, younger adults may have sufficient cognitive resources to adapt culturally-nonpreferred strategies or to overcome potential limitations of culturally-preferred strategies. Older adults, in contrast, may lack the cognitive resources to adapt or overcome limitations of their well-practiced culturally-supported strategies (Na, Huang, & Park, 2017; Park et al., 1999). For tasks that require high levels of cognitive resources, age-related declines in cognition could lead to pervasive and consistent effects of age across cultures. Aligning with this, discriminating old from new objects in our study is less cognitively demanding than discriminating old from similar lures. Thus, the presence of an interaction of age and culture in the old/new comparison may be in line with it being an easier task than discriminating old items from similar lures in which age effects seem to surpass cultural effects. In accordance with the patterns of behavioral findings, the neural findings appear to indirectly support the potential moderating role of cognitive load in the relationship between age and culture on memory (Gutchess & Cho, 2024; Na et al., 2017; Park et al., 1999). Specifically, cultural differences in brain regions related to higher-order cognitive processes (i.e., LIFG) were eliminated with age. In addition, simple age differences (rather than interactions of age and culture) were more consistently found for pattern separation than for old/new discrimination, possibly reflecting more cognitively demanding memory processes (Adams et al., 2022; Leal et al., 2017; Stark et al., 2010, 2013; Yassa, Mattfeld, Stark, & Stark, 2011). Approaches that target the cognitive demands of the task or probe individual differences in cognitive resources are crucial steps to further understand the ways in which culture affects age differences in memory and neural recruitment. Such an approach may help to better interpret the patterns of LIFG activity, as well as differences in the findings in the comparisons of old/new discrimination and pattern separation in the present study.

The current study, of course, has several limitations. Typical challenges to the study of aging are reflected here, including limitations of generalizability. Our samples of older adults are likely highly select, a concern that is exacerbated by the screening requirements to participate in an fMRI study. In addition, it is challenging to isolate effects of age in cross-sectional comparisons of younger and older adults. To address this concern, future studies should employ longitudinal designs or include middle-aged samples (Lachman, 2015). The effects of age observed in this study could, to some extent, reflect the contribution of cohort differences. This could also apply to cultural differences, should they reflect unintended differences between our samples, although we note how well our samples are across cultures on neuropsychological test performance. In addition, our analytic choices for the neuroimaging data likely affected our results. Given the novelty of cross-cultural investigations of pattern separation, as well as memory processes more generally, we adopted an approach that focused on regions previously implicated in cultural differences in memory in younger adults. Although we attempted to balance a targeted approach using ROIs to increase sensitivity to detect effects alongside exploratory whole-brain analyses, an omnibus ANOVA approach may not be sensitive enough to detect the effects of age and culture. The lack of effects in the hippocampus for pattern separation is surprising given that this is one of the key regions for discriminating old from similar lures (Baker et al., 2016; Bakker et al., 2008; Yassa & Stark, 2011) and strongly impacted by aging (Leal et al., 2017; Stark et al., 2010, 2013, 2015). Our lack of findings may reflect the use of hippocampal ROIs rather than investigating subregions of hippocampus (i.e., DG/CA3, CA1), which are typically probed using high-resolution imaging. However, recent research, including our own findings implicating LIFG, suggests that regions beyond the hippocampus related to higher-order cognitive processes (e.g., dorsal medial prefrontal cortex) can be involved in pattern separation (Nash, Hodges, Muncy, & Kirwan, 2021; Pidgeon & Morcom, 2016).

Despite the limitations, the present study contributes to our understanding of age, culture, and memory. To our knowledge, this is the first study examining the effects of age and culture on memory with not only behavioral but neural measures. Although several studies have found cross-cultural differences in memory specificity (Leger & Gutchess, 2021; Leger et al., 2024b; Mickley Steinmetz et al., 2018; Millar et al., 2013; Paige, Ksander, Johndro, & Gutchess, 2017), this is the first to test this in older adults. In summary, the results suggest that comparing the discrimination of old vs. new information at retrieval is sensitive to the influences of age and culture. Taiwanese performed worse than Americans, and the effects of age were more pronounced for the Taiwanese. Americans activated the hippocampus for new more than old items, but pattern of activity for the conditions did not differ for Taiwanese, nor did it interact with age. The engagement of LIFG differed across cultures such that the pattern of greater activity for old (for Americans) or new (for Taiwanese) items was lost with age, particularly for older Americans. We speculate that the results could reflect cultural differences in the orientation to novelty vs. familiarity for younger adults, with the LIFG engaged to support interference resolution at retrieval. Older adults, in contrast, lack the cognitive resources to successfully recruit LIFG, which may affect the engagement of this region for Americans more than Taiwanese. Support is not as strong for cultural differences in processes related to pattern separation. Although Americans had higher levels of memory discrimination than Taiwanese and engaged the LIFG for correct rejections more than false alarms, the patterns of behavior and neural activity did not interact with culture and age. Furthermore, we did not detect effects of culture or age in the hippocampus, the region most implicated in pattern separation. The findings suggest ways in which cultural life experiences and associated information processing strategies may contribute to consistent effects of age across cultures or lead to different trajectories with age in terms of memory performance and neural engagement.

## Acknowledgements

The authors thank Danielle Schwartz, Abaigeal Ford, Erin Wong, Nicolette Barber, Chi-Chuan Chen, Chun-Yi Lee, Li-An Her, Jennifer Jing-Yu Chuang, Yi-Hsiu Lee, and Hannah Lin-Han Huang for assistance with data collection and scoring. We are grateful to Yu-Ling Chang for helpful discussion and recommendations.

## Open Practices Statement

None of the data or materials for the experiments reported here is available, and none of the experiments was preregistered.

## Footnotes

1 A cultural comparison of young adults was published in Leger et al. (2024a). However, the young adult samples between Leger et al. (2024a) and the current paper differ in that one Taiwanese young adult from the previous paper was excluded in the present study due to the use of different criteria to determine excessive motion (please see the “data acquisition and preprocessing” section for details). In addition, behavioral data were recalculated to exclude trials with missing responses, rather than treating those trials as errors.

2 For analyses, 4 American older adults only had data from 3 runs (out of 4) due to experimenter error or the participant terminating the scan mid-run. In cases where parts of runs were repeated, we used the trials in which participants saw the images for the first time.

3 Comparison of young adult resting state data across cultures has been published in Zhang et al. (2022).

4 Directly comparing d’s for young Americans and Taiwanese on the target-foil condition yields a significant cultural difference, *p* = .02

